# Marine Picoplankton Metagenomes from Eleven Vertical Profiles Obtained by the Malaspina Expedition in the Tropical and Subtropical Oceans

**DOI:** 10.1101/2023.02.06.526790

**Authors:** Pablo Sánchez, Marta Sebastián, Massimo Pernice, Raquel Rodríguez-Martínez, Stephane Pesant, Susana Agustí, Takashi Gojobori, Ramiro Logares, María Montserrat Sala, Dolors Vaqué, Ramon Massana, Carlos M. Duarte, Silvia G. Acinas, Josep M. Gasol

## Abstract

The Ocean microbiome has a crucial role on Earth’s biogeochemical cycles, but also represents a tremendous potential for biological applications as part of the blue-biotechnology. During the last decade, global cruises such as *Tara* Oceans or the Malaspina Expedition have expanded our knowledge on the diversity and genetic repertoire of marine microbes. Nevertheless, there is still a gap of knowledge on broad scale patterns between photic and bathypelagic dark ocean microbes derived from the lack of detailed vertical profiles covering contrasting oceans depth regions. Here we present a dataset of 76 microbial metagenomes of the picoplankton size fraction (0.2-3.0 μm) collected in 11 stations along the Malaspina Expedition circumnavigation that cover vertical profiles sampling at 7 depths, from the surface to the 4000 m deep (or the sea floor in shallower waters). This Malaspina Microbial Vertical Profiles metagenomes (MProfile) dataset produced 1.66 Tbp of raw DNA sequences that assembled into a total 25.3 Gbp. After gene prediction and annotation, we built a 46.3 million non-redundant gene compendium with their corresponding annotations (M-GeneDB-VP), clustered at 95% sequence similarity. This dataset will be a valuable resource for exploring the functional and taxonomic connectivity between the photic and bathypelagic tropical and subtropical ocean at a global scale, while increasing our general knowledge on the Ocean microbiome.

## BACKGROUND AND SUMMARY

The ocean is the largest biome on Earth. Most of its biomass and biodiversity is due to microorganisms, mainly bacteria and archaea ^1^, that have a crucial role in biogeochemical cycles ^2^. After pioneering work in the GOS expedition ^3^, the main worldwide exploration of the marine microbiome through the analyses of microbial metagenomes have been those of the *Tara* Oceans Expedition (2009-2013) ^4^, the Malaspina 2010 Expedition ^5^, and more recently the Bio-GEOTRACES ^6^ and Bio-GO-SHIP programs ^7^.

Specifically, the Malaspina 2010 Circumnavigation Expedition ^5^ sampled the marine microbiome in tropical and sub-tropical oceans, from the surface down to bathypelagic waters (~4,000 m depth) between 2010 and 2011. This expedition has contributed to the exploration of the patterns of microbial diversity and biogeography both in the sunlit and dark oceans. In the photic ocean, the analyses of prokaryotic 16S TAGs metabarcoding datasets pointed to shifts towards communities enriched in rare taxa reflecting environmental transitions ^8^ and both 16S and 18S TAGs metabarcoding highlighted the role of dispersion on planktonic and micro-nektonic organisms ^9^. Also, the patterns in viral abundance, production and the role of viral lysis as a driver of prokaryote mortality have been assessed from surface to bathypelagic layers ^10^. In the dark ocean, the Malaspina expedition contributed with a survey of the diversity and biogeography of deep-sea pelagic prokaryotes ^11^ as well as that of heterotrophic protists, unveiling the special relevance of fungal taxa ^12^. It also determined that the particle-association lifestyle is a phylogenetically conserved trait in bathypelagic prokaryotes ^13^. The first 58 microbial metagenomes of the bathypelagic ocean allowed us to reconstruct 317 high-quality metagenome-assembled genomes to metabolically characterize the deep ocean microbiome ^14^. However, the Malaspina *-omics* datasets have not yet been fully exploited. Now, we present a new metagenomic resource of the ocean picoplankton, with particular emphasis on the description of detailed vertical profiles, from surface photic layers down to 4,000 m deep, covering the deep chlorophyll maximum (DCM), the mesopelagic and the bathypelagic realm with 3-4 depths. Therefore, the Malaspina Microbial Vertical Profiles metagenomes dataset (MProfile) complements previous metagenomic data sets derived from *Tara* Oceans expedition, that sampled surface waters through the mesopelagic ocean ^15^ and our previous Malaspina bathypelagic deep ocean metagenomes dataset ^14^.

This resource consists on:

i. primary data in the form of 1.66 Tbp of environmental whole genome shotgun sequencing data, distributed in 76 samples (Fig. 1) corresponding to 7 depths in 11 vertical profiles (108.13 ± 2.831 million read pairs, mean ± sd, and 21.84 ± 0.572 Gbp per sample), along the R/V Hespérides track across the global tropical and sub-tropical ocean during 2010-2011.
ii. a total of 25.3 Gbp of assembled contigs (332.9 Mbp ± 50.25 per sample, mean ± standard deviation; Supplementary Table 4). A fraction of 67.07% ± 1.84 of the predicted coding DNA sequences (CDS) in the assembled contigs (510,378 ± 87,562.5) could be assigned to at least one functional category (Fig. 3): 31.3% ± 1.53 to clusters of orthologous groups (COG)^16^, 62.0% ± 1.86 to protein families (PFAM)^17^, 26.7% ± 1.89 to Kyoto Encyclopedia of Genes and Genomes (KEGG)^18^ orthologs (KO), and 1.1% ± 0.13 to carbohydrate-active enzymes (CAZy)^19^. A fraction of 36.93% ± 1.846 of CDS could not be assigned to a function with the used databases (Supplementary Table 4). Similarly, 41.7% ± 10.7 CDS (207,270 ± 47,542.0) per sample could not be taxonomically classified further than “root” or were “unclassified” after aligning them to UniRef90 ^20^ using the lowest common ancestor approach (LCA).
iii. a 46.3 million non-redundant CDS compendium, the Malaspina Vertical Profiles Gene Database (M-GeneDB-VP). In total 24,107,535 (52.1%) genes in this gene catalog were annotated with PFAM, 9,013,490 (19.48%) with KOs, 10,170,434 (22.0%) with COGs and 467,469 (1.01%) with CAZy, whereas 21,754,323 genes (47.0%) could not be annotated and correspond to the gene novelty of this database.
iv. functional profiles of each gene grouped by annotation, consisting on the abundance of each CDS/annotation per sample, based on the number of reads of each metagenome mapping back to the M-GeneDB-VP, corrected by gene length and by single-copy universal marker gene abundance.
v. taxonomic profiles of picoplankton, based on the 16S and 18S mTAG ^21^ analysis of the metagenomes, including more than 15,000 OTUs (Fig. 4).

**Figure 1.**
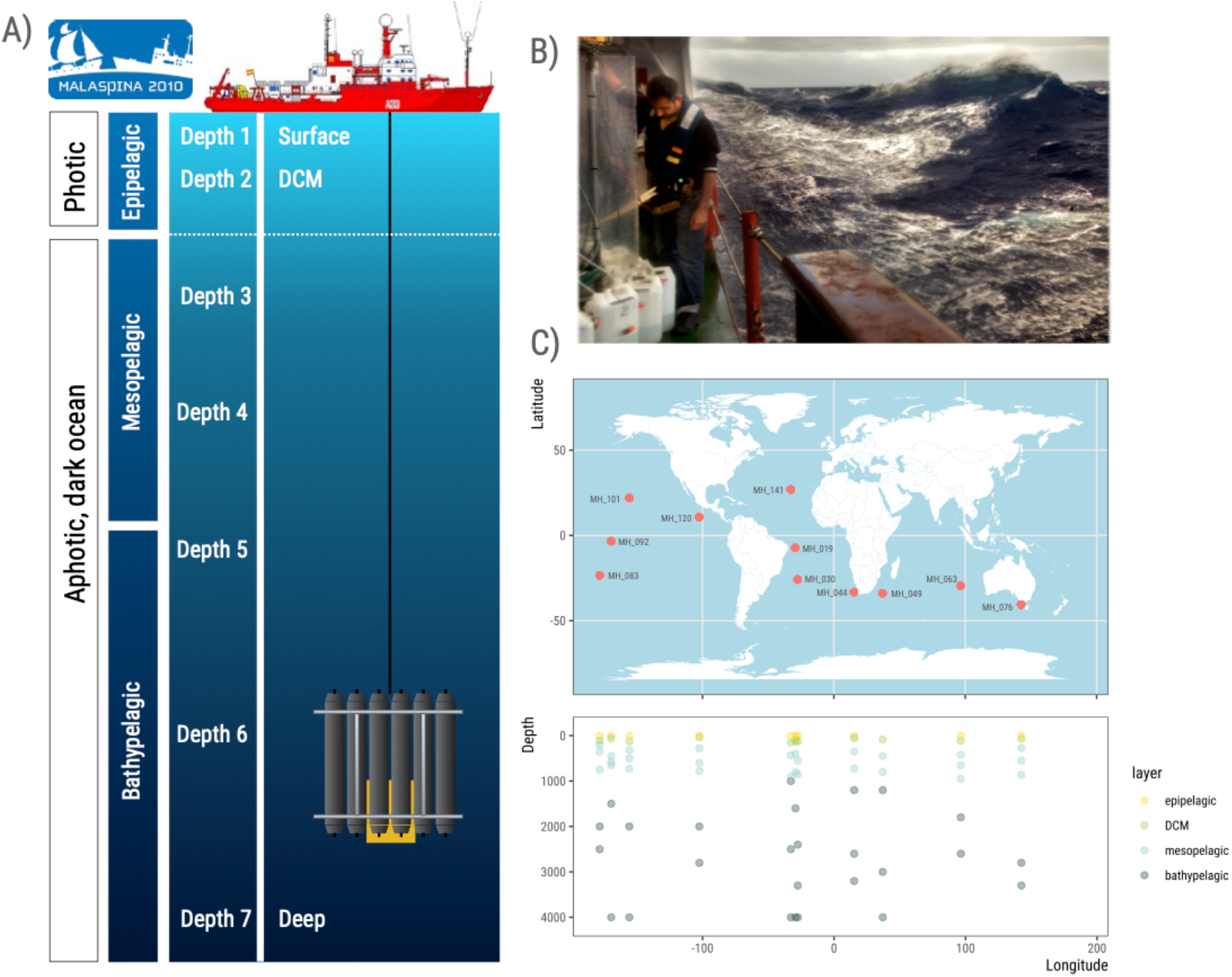
The Malaspina expedition prokaryotic vertical profiles. **A)** Schematics of a typical vertical profile sampling event. Water samples at seven depths including deep chlorophyll maximum (DCM), the deep scattering layer (DSL) and a deep sample (close to the sea floor or 4,000 m deep), targeting 3 layers from the photic and dark ocean: epipelagic, including DCM, mesopelagic and bathypelagic. **B)** Dr. Massimo Ciro Pernice sampling the CTD-rosette in the South Pacific Ocean. Photo credit: Joan Costa/CSIC. **C)** Map showing the sampling stations of the Malaspina Expedition presented in this data set, along the tropical and sub-tropical global Ocean, and the depths from where water was collected for whole genome shotgun sequencing of the 0.2 μm-3 μm size fraction.

**Figure 2.**
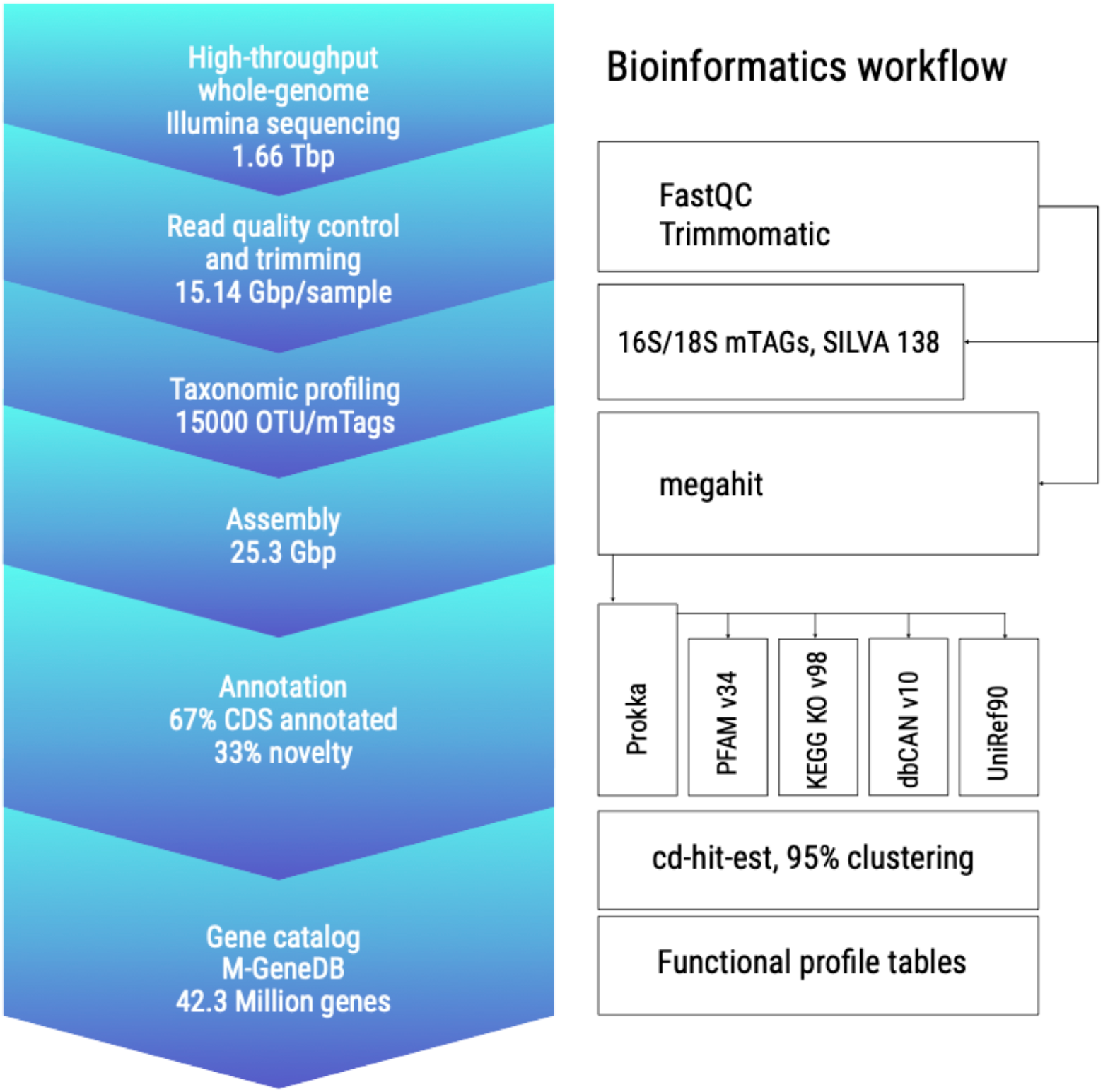
Bioinformatics workflow. Summary of the different steps taken to process this dataset, from sequencing to the final gene catalog and taxonomic and functional profiles, highlighting some of the metrics obtained in each part of the workflow, side by side to some of the bioinformatics tools used in the analysis.

**Figure 3.**
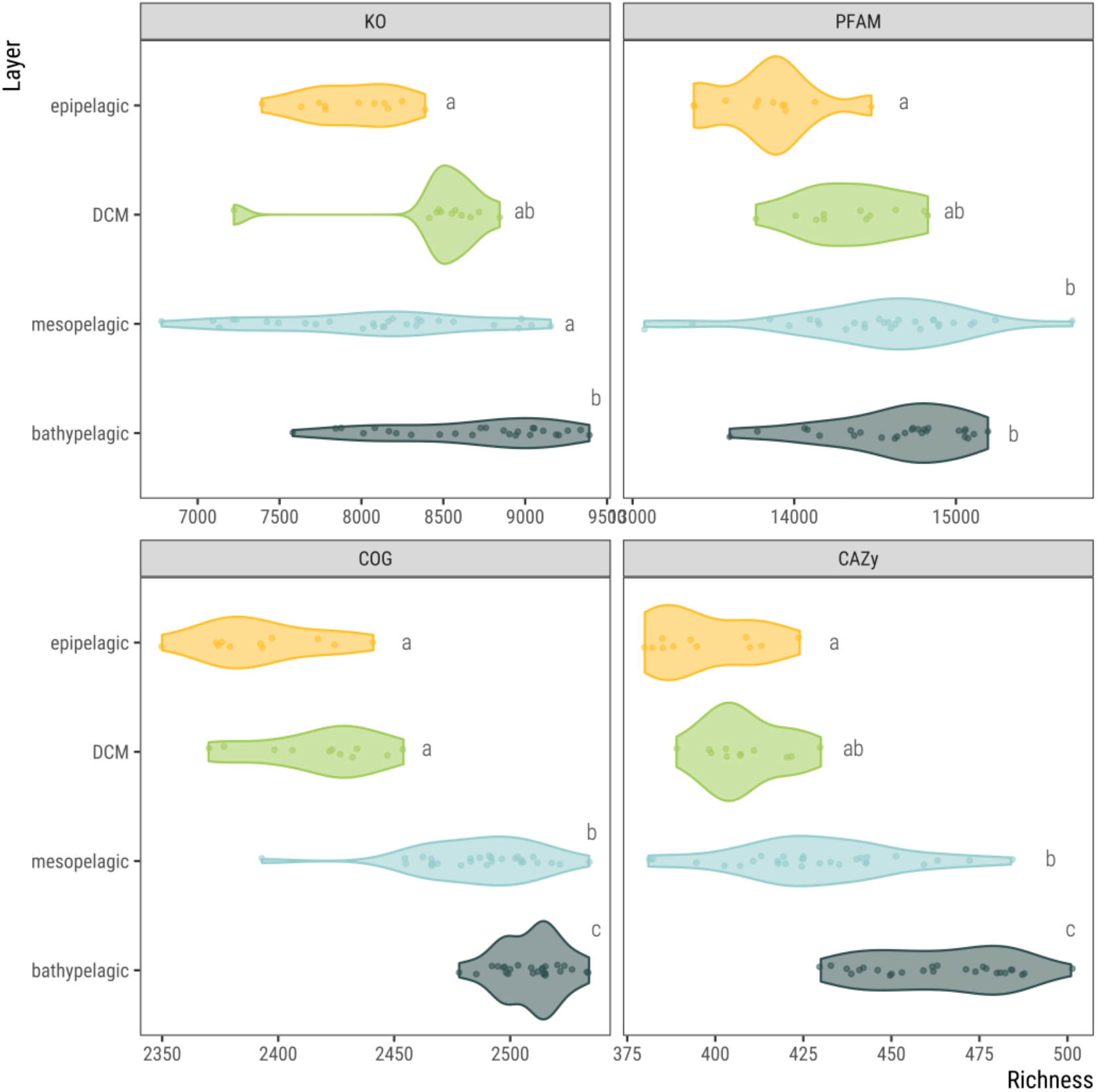
Functional richness. Functional richness of 76 metagenomes from 11 vertical profiles from the Malaspina Expedition showed by ocean layer: epipelagic (from 0 to 200m deep), including the deep chlorophyll maximum (DCM), mesopelagic (200 to 1,000m deep) and bathypelagic (1,000 to 4,000m deep). Richness is calculated as the number of distinct annotations (KEGG orthologs; KO, protein families; PFAM, clusters of orthologous groups; COG and carbohydrate-active enzymes; CAZy) in each sample. Significant differences in richness values between ocean layers are depicted with different letters (Kruskal-Wallis, p<0.05; Dunn’s post-hoc test with Holm correction for multiple comparisons).

**Figure 4.**
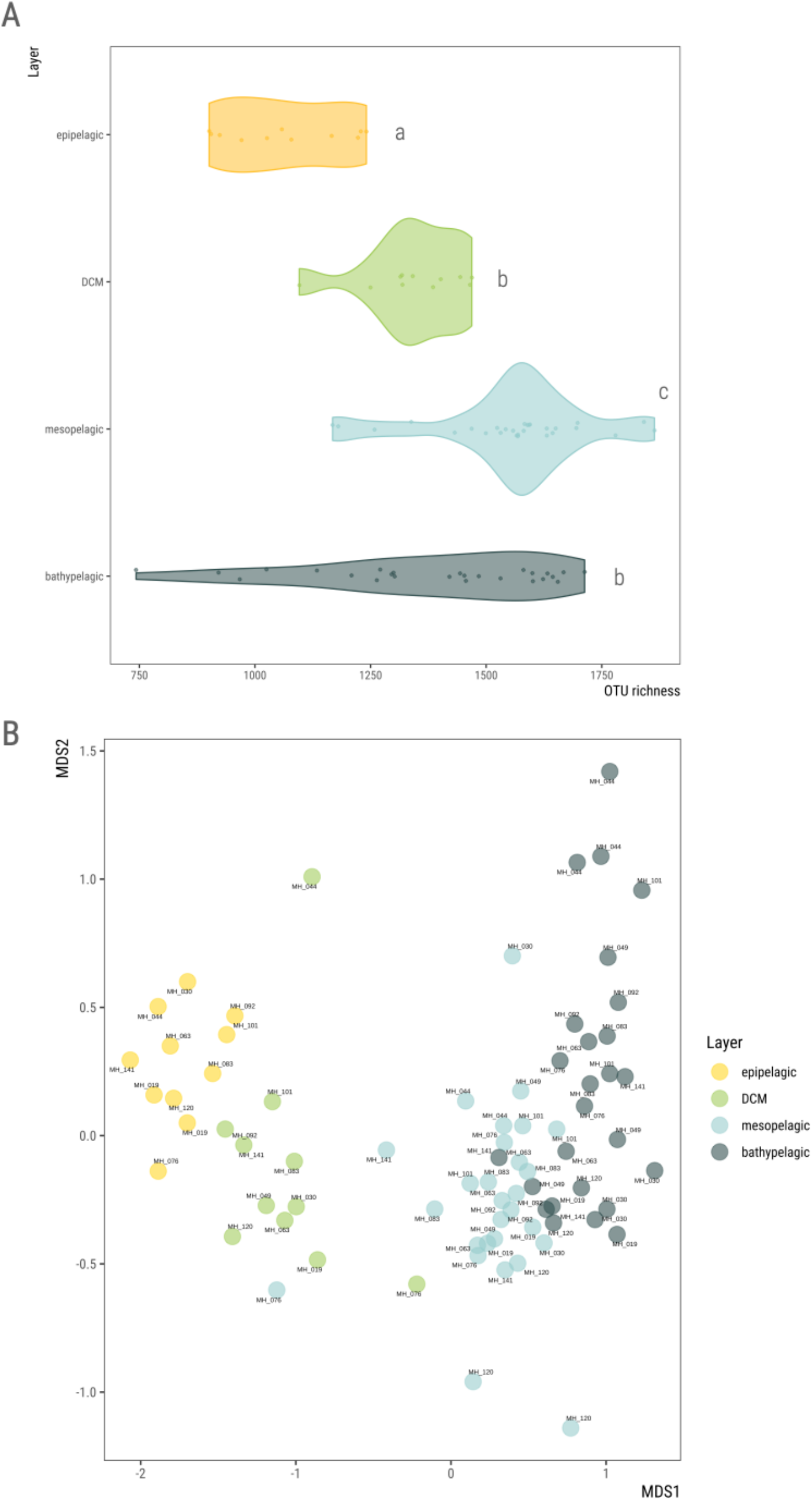
16S and 18S mTAGs taxonomic profiling. **A)** OTU richness based on mTAGs (16S and 18S rRNA) analysis, showed by ocean layer: epipelagic (from 0 to 200m deep), including the deep chlorophyll maximum (DCM), mesopelagic (200 to 1,000m deep) and bathypelagic (1,000 to 4,000m deep). Significant differences in richness values between ocean layers are depicted with different letters (Kruskal-Wallis, p<0.05; Dunn’s post-hoc test with Holm correction for multiple comparisons). **B)** Ordination plot (non-metric multidimensional scaling; Bray-Curtis distance) of 76 metagenomes from 11 vertical profiles based on their community composition (mTAG OTUs) colored by ocean layer as described above.

This Malaspina Microbial Vertical Profiles metagenomes dataset resource will be of great interest to the community to tackle ecologically relevant questions on marine microbial ecology, such as inferring differential functional traits of photic and aphotic bacterial and archaeal genomes through the reconstruction of metagenome assembled genomes (MAGs) ^22^, serving as a valuable dataset for gene discovery with interest in biotechnology, such as the finding of new CRISPR-Cas systems in the deep ocean ^23^ and other research areas.

Primary sequencing data has been submitted to the European Nucleotide Archive and derived data such as M-GeneDB-VP, assemblies, annotations and files for functional and taxonomic profiles will be soon submitted to public repositories (BioStudies), to allow further exploration of the functional and taxonomic composition and vertical connectivity of the ocean microbiome.

## METHODS

### 1. Sample collection

A total of 76 water samples were taken during the Malaspina 2010 expedition (http://www.expedicionmalaspina.es) on board the R/V Hespérides, corresponding to 11 different sampling stations (Fig. 1a). Each station was profiled by collecting water from up to 7 discrete depths from surface (3 m) to the bathypelagic layer, down to 4,000 m (mean maximum depth of each profile 3,491 ± 626.8 m, ± standard deviation), including the DCM (Fig. 1c; Supplementary Table 2). Water samples were collected either with a rosette of Niskin bottles (12 L each) on a frame with a CTD sensor or with a large Niskin bottle (30 L) for the surface samples. For every sample, two replicates with a volume of 6 L of the sampled material were pre-filtered sequentially on a 200 μm and 20 μm nylon mesh to remove large plankton, and then through a 47 mm diameter polycarbonate (PC) membrane with a 3 μm pore size (Whatman filter ref: 10418312), and a 47 mm diameter PC membrane with a 0.22 μm pore size (Whatman filter ref:GTTP04700) with a peristaltic pump (Masterflex, EW-7741010) with a flow rate of 50-100 ml min^-1^. When the filtration rate decreased considerably, filters were replaced. The 0.22 μm filters, enriched in biomass of the free-living (FL) prokaryotic community ^24,25^, were packaged in 2-mL cryotubes, treated with no addition of chemical, labelled with a barcode identification sticker, flash frozen in liquid nitrogen and stored in a freezer at −80 °C. The two replicate filters for each of the samples were stored in 2 distinct tubes with the same sample identifier. Latex or nitryl gloves were used for this protocol. All containers, filter holders and tubing were washed with 0.1% bleach and rinsed with milliQ water. All tweezers were kept clean with ethanol. The time span from bottle closing of the deep sample to filter freezing was around 4 h and except for the time needed to empty the rosette bottles, the water was kept at 4 °C.

### 2. DNA extraction

DNA was extracted with the standard phenol-chloroform protocol with slight modifications ^21,26^. Detailed description of the DNA extraction protocol used in our lab has been published before ^27^. Briefly, the filters were cut in small pieces with sterile razor blades and half of each filter was resuspended in 3 ml of lysis buffer (40 mM EDTA, 50 mM Tris-HCl, 0.75 M sucrose). Samples were incubated at 37 °C for 45 min in lysis buffer (Lysozyme; 1 mg ml^-1^ final concentration) with gentle agitation. Then, the buffer was supplemented with sodium dodecyl sulfate (SDS, 1% final concentration) and proteinase K (0.2 mg ml^-1^ final concentration) and the samples were incubated at 55 °C for 60 min under gentle agitation. The lysate was collected and processed with the standard phenolchloroform extraction procedure: an equal volume of Phenol:CHCl3:IAA (25:24:1, vol:vol:vol) was added to the lysate, mixed and centrifuged 10 min at 3000 rpm. Then the aqueous phase was recovered and the procedure was repeated. Finally, residual phenol was removed by adding an equal volume of CHCl3:IAA (24:1, vol:vol) to the recovered aqueous phase. The mixture was centrifuged and the aqueous phase was recovered for further purification. The aqueous phase was then concentrated by centrifugation with a Centricon concentrator (Millipore, Amicon Ultra-4 Centrifugal Filter Unit with Ultracel-100 membrane). This step was repeated three times by adding 2 ml of sterile MilliQ water each time in order to increase the DNA purity up to a target of 100 to 200 μl of total genomic DNA.

### 3. Sequencing

Extracted DNA was sequenced on the 2×101 bp Illumina HiSeq2000 platform on the Centre Nacional d’Anàlisi Genòmica (CNAG) in Barcelona, Spain, yielding a total of 1.66 Tbp (108.13 ± 2.831 million read pairs and 21.84 ± 0.572 Gbp per sample; mean ± standard deviation). Fastq files with the clean reads for all 76 samples are available at ENA under the BioProject accession number PRJEB52452 (Supplementary Table 1).

### 4. Bioinformatics workflow

The bioinformatics workflow applied to this data set is summarized in Fig. 2.

#### 4.1. Quality control and raw read trimming

The quality of raw read pairs was checked with fastqc v0.11.7 (https://www.bioinformatics.babraham.ac.uk/projects/fastqc/), and Illumina TruSeq adapter contamination was removed in trimmomatic v0.38 ^28^ keeping adapter-free read pairs with contiguous quality over 20 and a minimum length of 45 bp with options *“ILLUMINACLIP:2:30:10 LEADING:3 SLIDINGWINDOW:4:20 MINLEN:45”*. Unpaired reads were discarded for further steps. After trimming the dataset consisted on a total of 1.15 Tbp, with 81.53 ± 4.553 million clean read pairs and 15.14 ± 1.049 Gbp per sample (Supplementary Table 3).

#### 4.2. Assembly, gene prediction and annotation

Clean reads were assembled in megahit v1.1.3 ^29^ with options *“--presets meta-large-- min-contig-len 500”* to produce 25.3 Gbp of metagenomic assemblies (332.9 Mbp ± 50.25). Assembled contigs larger than 500 bp were annotated in prokka v1.14.6 ^30^ for gene prediction based in prodigal ^31^, COGs ^16^, Enzyme Commission numbers (EC) and gene product name. Additionally, predicted genes amino acid sequences were annotated for protein families’ domains (PFAM v34) ^17^ using HMMER v3.33 *(hmmsearch)*^32^ with option *“-E 0.1”,* KEGG KOs ^18^ release v98.0 using kofamscan v1.3.0 ^33^ and options *format detail-E 0.01”,* and CAZymes using HMMER v3.33 *(hmmsearctì)* against dbCAN v10 ^19^.

PFAM *hmmsearch* results were obtained with very low stringency (E=0.1), so the best hit was awarded for each model that aligned with no overlap to the predicted genes. This means that a single gene might have more than one protein domain annotation. When two or more hits were aligning in the same region, if the overlap was longer than half the length of the smaller alignment, only the hit bit larger bitscore was retained for that region.

Similarly, kofamscan (E=0.01) results were filtered by keeping all hits with scores above the predefined thresholds for individual KOs (marked with an ‘*’), potentially assigning more than one KO to a single predicted gene.

Each predicted coding sequence was taxonomically assigned by mapping them to UniRef90 ^20^, release 2021_03 from 9 of June 2021, with mmseqs2 ^34^ development version, commit 13-45111, with the taxonomy workflow options *“--max-accept 100 -tax-lineage 1 -e 1E-5 -v 3 -a*” and converted to table with *mmseqs createtsv*. All ranks out of domain, phylum, class, order, family, genus or species were removed from classification and missing fields were marked as “unclassified”. The lowest common ancestor for each sequence was also recorded.

#### 4.3. Gene catalog

In order to condense the genetic variability of the predicted gene dataset, we clustered all coding DNA sequences longer than 100 bp to 95% nucleotide sequence similarity and 90% alignment coverage of the shorter sequence in cd-hit-est v4.6.1 ^35^ with options “*-c 0.95 -G 0 -aS 0.9 -g 1 -r 1 -d 0 -s 0.8”*. We used the longest sequence of each cluster as the representative sequence, obtaining a catalog of 46,279,067 non-redundant genes, referred to as the Malaspina Vertical Profiles Gene Database (M-GeneDB-VP). Functional and taxonomic annotation of the M-GeneDB-VP genes was inherited from the annotation of representative sequence of each cluster, as described above.

#### 4.4. Functional profiling

Clean reads were back-mapped to the catalog with bowtie2 v2 2.2.9 ^36^ and alignments were filtered with samtools v1.3.1 ^37^ with options *“-q 10 -F 260”* to keep only primary alignments. Reads mapping to catalog genes were counted in htseq-count v0.10.0 ^38^ with options *“--nonunique all”* to build gene profiles per sample. As genes in a catalog are stripped from their genomic context, this strategy allowed us to count a read mapping to all features it was assigned to, instead of randomly imputing it to only one. Counts were normalized by gene length in bp and then normalized by the geometric median abundance of 10 universal single-copy phylogenetic marker genes either for COGs (COG0012, COG0016, COG0018, COG0172, COG0215, COG0495, COG0525, COG0533, COG0541, and COG0552) or KOs (K01409, K01869, K01873, K01875, K01883, K01887, K01889, K03106, K03110, K06942) respectively. Normalizing coverage-corrected read counts by the abundance of these marker genes acts as a proxy to the number of gene copies per cell ^39^. Functional profiles for COGs, PFAMs, KOs and CAZymes were calculated by adding up abundance values corresponding to genes annotated as a particular function, both from the gene length normalized table and the single-copy marker gene normalized table. Functional richness was calculated by converting gene length normalized tables to pseudo-counts (multiplying abundance values by 10,000 and rounding to the next integer) and rarefying to 0.95 times the minimal sample sum with function *rtk* in R package rtk v0.2.6.1 ^40^.

#### 4.5. Taxonomic profiling

Taxonomic profiling of the samples was carried out by identifying and classifying mTAGs, ribosomal RNA gene small subunit (SSU) fragments directly from the Illumina-sequenced metagenomes (Logares et al., 2014) with mTAGs v1.0.4 ^41^, profile workflow with options *“-ma 1000 -mr 1000”.* This protocol is particularly suitable for metagenomes with short reads, as it takes advantage of a degenerated consensus reference database and an exhaustive search strategy, reducing the number of ambiguously mapped sequences that could not be used for classification. Briefly, the mTAGs pipeline extracts reads from the metagenome which are compatible with the SSU-rRNA gene sequence by using hidden Markov models, and then maps them to a sequence database based on SILVA 138 ^42^, pre-clustered at 97% of sequence similarity and with degenerated consensus sequences within each OTU. It then classifies mTAGs conservatively to a taxonomic rank by considering its lowest common ancestor. Finally, it builds taxonomic profiles at different ranks, including the OTU level. OTUs classified as “class Cyanobacteriia; order Chloroplast” were removed from the OTU counts table. The OTU count table was rarefied (5,861 reads/sample) using the *rrarefy* function in the R package vegan v2.5.7^43^ to correct for uneven sequencing depths among samples.

## DATA RECORDS

All sequencing products described here can be found under BioProject accession number PRJEB52452 hosted by the European Nucleotide Archive. ENA accession numbers for each metagenome sequencing runs are provided in Supplementary Table 1.

Derived products such as the metagenome assemblies, the M-GeneDB-VP gene catalog, or the functional and taxonomic profiles will be deposited in the European Bioinformatics Institute BioStudies database in a future publication.

Underway and meteorological data measured on board R/V Hesperides can be accessed at the Marine Technology Unit (UTM, CSIC) http://data.utm.csic.es/geonetwork/srv/eng/catalog.search.

## TECHNICAL VALIDATION

Extracted DNA was quantified using a Nanodrop ND-1000 spectrophotometer (NanoDrop Technologies Inc, Wilmington, DE, USA) and the Quant_iT dsDNA HS Assay Kit with a Qubit fluorometer (Life Technologies, Paisley, UK).

The sequencing error rate was calculated by the sequencing center using PhiX147 phage DNA spikes (0.44% ± 0.138).

## CODE AVAILABILITY

All the software used to process the data set presented here is publicly available and distributed by their developers. All versions have been specified in the main text, along with the options used departing from defaults. Custom scripts used in intermediate or summarizing steps are available at https://gitlab.com/malaspina-public/picoplankton-vertical-profiles.

## ACKNOWLEDGEMENTS

We thank the R/V Hespérides captain and crew, the chief scientists in the Malaspina expedition legs, and all project participants for their help in making this project possible. This work was funded by the Spanish Ministry of Economy and Competitiveness (MINECO) through the Consolider-Ingenio program (Malaspina 2010 Expedition, ref. CSD2008-00077). The sequencing of 76 metagenomes from 11 vertical profiles was funded by project Malaspinomics and Malaspina-analytics (CTM2011-15461-E) awarded to C.M.D. by the Spanish Ministry of Economy and Competitiveness. Additional funding was provided by the project MAGGY (CTM2017-87736-R) to S.G.A. from the Spanish Ministry of Economy and Competitiveness, Grup de Recerca 2017SGR/ 1568 from Generalitat de Catalunya, and King Abdullah University of Science and Technology (KAUST) under contract OSR #3362. The ICM researchers have had the institutional support of the “Severo Ochoa Centre of Excellence” accreditation (CEX2019-000928-S). High-Performance computing analyses were run at the Marine Bioinformatics Core facility (MARBITS, https://marbits.icm.csic.es) of the Institut de Ciències del Mar (ICM-CSIC), Barcelona Supercomputing Center (Grant BCV-20132-0001), KAUST’s Ibex HPC and Galicia Supercomputing Center (CESGA).

## AUTHOR’S CONTRIBUTIONS

P.S. analyzed the data, wrote custom code in perl, python, bash and R, and wrote the manuscript. M.S., M.C.P. and R.G. extracted DNA. M.C.P. also participated in the sampling operations. S.P and P.S. curated the metadata and defined the biosamples. S.A. contributed funding. T.G. contributed funding. C.M.D. was the chief coordinator of the Malaspina Expedition and contributed funding and computational assistance. S.G.A. defined sampling protocols, coordinated the microbial -*omics* analyses and contributed funding. J.M.G. coordinated the microbial diversity and ecosystem function area of the Malaspina Expedition and contributed funding.

## REFERENCES

1. Bar-On, Y. M., Phillips, R. & Milo, R. The biomass distribution on Earth. Proc. Natl. Acad. Sci. 115, 6506–6511 (2018).

2. Cho, B. C. & Azam, F. Major role of bacteria in biogeochemical fluxes in the ocean’s interior. Nature 332, 441–443 (1988).

3. Yooseph, S. et al. The Sorcerer II global ocean sampling expedition: Expanding the universe of protein families. PLoS Biol. 5, e16 (2007).

4. Karsenti, E. et al. A holistic approach to marine Eco-systems biology. PLoS Biol. 9, e1001177 (2011).

5. Duarte, C. M. Seafaring in the 21St Century: The Malaspina 2010 Circumnavigation Expedition. Limnol. Oceanogr. Bull. 24, 11–14 (2015).

6. Biller, S. J. et al. Marine microbial metagenomes sampled across space and time. Sci. Data 5, 180176 (2018).

7. Larkin, A. A. et al. High spatial resolution global ocean metagenomes from Bio-GO-SHIP repeat hydrography transects. bioRxiv 2020.09.06.285056 (2020)

8. Ruiz-González, C. et al. Higher contribution of globally rare bacterial taxa reflects environmental transitions across the surface ocean. Mol. Ecol. 28, 1930–1945 (2019).

9. Villarino, E. et al. Large-scale ocean connectivity and planktonic body size. Nat. Commun. 9, 142 (2018).

10. Lara, E. et al. Unveiling the role and life strategies of viruses from the surface to the dark ocean. Sci. Adv. 3, e1602565 (2017).

11. Salazar, G. et al. Global diversity and biogeography of deep-sea pelagic prokaryotes. ISME J. 10, 596–608 (2016).

12. Pernice, M. C. et al. Global abundance of planktonic heterotrophic protists in the deep ocean. ISME J. 9, 782–792 (2015).

13. Salazar, G. et al. Particle-association lifestyle is a phylogenetically conserved trait in bathypelagic prokaryotes. Mol. Ecol. 24, 5692–5706 (2015).

14. Acinas, S. G. et al. Deep ocean metagenomes provide insight into the metabolic architecture of bathypelagic microbial communities. Commun. Biol. 4, 1–15 (2021).

15. Sunagawa, S. et al. Structure and function of the global ocean microbiome. Science 348, 1261359 (2015).

16. Galperin, M. Y., Makarova, K. S., Wolf, Y. I. & Koonin, E. V. Expanded Microbial genome coverage and improved protein family annotation in the COG database. Nucleic Acids Res. 43, D261–D269 (2015).

17. El-Gebali, S. et al. The Pfam protein families database in 2019. Nucleic Acids Res. 47, D427–D432 (2019).

18. Kanehisa, M. & Goto, S. KEGG: Kyoto Encyclopedia of Genes and Genomes. Nucleic Acids Res. 28, 27–30 (2000).

19. Yin, Y. et al. dbCAN: a web resource for automated carbohydrate-active enzyme annotation. Nucleic Acids Res. 40, W445–51 (2012).

20. Suzek, B. E., Wang, Y., Huang, H., McGarvey, P. B. & Wu, C. H. UniRef clusters: A comprehensive and scalable alternative for improving sequence similarity searches. Bioinformatics 31, 926–932 (2015).

21. Logares, R. et al. Metagenomic 16S rDNA Illumina tags are a powerful alternative to amplicon sequencing to explore diversity and structure of microbial communities. Environ. Microbiol. 16, 2659–2671 (2013).

22. Coutinho, F. H., Sánchez, P., Duarte, C. M., Gasol, J. M. & Acinas, S. G. Differential Functional traits between photic and aphotic prokaryotic genomes. Prep.

23. Sánchez, P. et al. Novel CRISPR-Cas systems at the temperate and tropical global deep-ocean. Prep.

24. Crump, B. C., Armbrust, E. V. & Baross, J. A. Phylogenetic Analysis of Particle–Attached and Free-Living Bacterial Communities in the Columbia River, Its Estuary, and the Adjacent Coastal Ocean. Appl. Environ. Microbiol. 65, 3192–3204 (1999).

25. Ghiglione, J. F., Conan, P. & Pujo-Pay, M. Diversity of total and active free-living vs. particle-attached bacteria in the euphotic zone of the NW Mediterranean Sea. FEMS Microbiol. Lett. 299, 9–21 (2009).

26. Mestre, M. et al. Sinking particles promote vertical connectivity in the ocean microbiome. Proc. Natl. Acad. Sci. 115, E6799–E6807 (2018).

27. Salazar, G. et al. Global diversity and biogeography of deep-sea pelagic prokaryotes. ISME J. 10, 1–13 (2015).

28. Bolger, A. M., Lohse, M. & Usadel, B. Trimmomatic: a flexible trimmer for Illumina sequence data. Bioinforma. Oxf. Engl. 30, 2114–2120 (2014).

29. Li, D. et al. MEGAHIT v1.0: A fast and scalable metagenome assembler driven by advanced methodologies and community practices. Methods 102, 3–11 (2016).

30. Seemann, T. Prokka: Rapid prokaryotic genome annotation. Bioinformatics (2014) doi:10.1093/bioinformatics/btu153.

31. Hyatt, D. et al. Prodigal: prokaryotic gene recognition and translation initiation site identification. BMC Bioinformatics 11, 119 (2010).

32. Eddy, S. R. Accelerated profile HMM searches. PLoS Comput. Biol. 7, e1002195, 1–16 (2011).

33. Aramaki, T. et al. KofamKOALA: KEGG ortholog assignment based on profile HMM and adaptive score threshold. Bioinformatics 1–2 (2019) doi:10.1093/bioinformatics/btz859.

34. Steinegger, M. & Söding, J. MMseqs2 enables sensitive protein sequence searching for the analysis of massive data sets. Nat. Biotechnol. 35, 1026–1028 (2017).

35. Li, W. & Godzik, A. Cd-hit: A fast program for clustering and comparing large sets of protein or nucleotide sequences. Bioinformatics 22, 1658–1659 (2006).

36. Langmead, B. & Salzberg, S. L. Fast gapped-read alignment with Bowtie 2. Nat. Methods 9, 357–359 (2012).

37. Li, H. et al. The Sequence Alignment/Map format and SAMtools. Bioinformatics 25, 2078–2079 (2009).

38. Anders, S., Pyl, P. T. & Huber, W. HTSeq--a Python framework to work with high-throughput sequencing data. Bioinforma. Oxf. Engl. 31, 166–169 (2015).

39. Salazar, G. et al. Gene Expression Changes and Community Turnover Differentially Shape the Global Ocean Metatranscriptome. Cell 179, 1068–1083.e21 (2019).

40. Saary, P., Forslund, K., Bork, P. & Hildebrand, F. RTK: efficient rarefaction analysis of large datasets. Bioinformatics 33, 2594–2595 (2017).

41. Salazar, G., Ruscheweyh, H.-J., Hildebrand, F., Acinas, S. G. & Sunagawa, S. mTAGs: taxonomic profiling using degenerate consensus reference sequences of ribosomal RNA genes. Bioinformatics 38, 270–272 (2022).

42. Quast, C. et al. The SILVA ribosomal RNA gene database project: improved data processing and web-based tools. Nucleic Acids Res. 41, D590–D596 (2013).

43. Dixon, P. VEGAN, a package of R functions for community ecology. J. Veg. Sci. 14, 927–930 (2003).

